# Phage T5 two-step injection

**DOI:** 10.1101/866236

**Authors:** John Davison

## Abstract

*Escherichia coli* bacteriophage T5 differs from most phages in that it injects its genome in two steps: First Step Transfer, FST, corresponding to leftmost 7.9% of the genome and Second Step Transfer, SST, corresponding to the remainder of the genome. Expression of genes *A1* and *A2* is required for SST. DNA injection stops at a site known as the injection stop signal (iss) which is a cis acting site located in the large untranslated region of the Left Terminal Repeat (LTR). The *iss* region is extremely complicated with many repeats, inverted repeats and palindromes. This report compares the *iss* regions of 25 T5-like phages and shows that all have a common conserved structure including a series of 8 DnaA boxes arranged in a highly specific manner; reminiscent of the origin of replication *(oriC)* of *E. coli.* DnaA protein, which binds to DnaA boxes, is a mostly membrane bound. A new, radically different, mechanism of T5 2-step injection is proposed whereby injecting T5 DNA stops at the plasma membrane due to the binding of the *iss* DnaA boxes to membrane-bound DnaA protein. Injection of the SST continues later via the combined action of the A1 and A2 proteins which cleave the FST DNA at a site upstream (right) of the *iss* region, thereby liberating it. They also cleave the incoming SST DNA at the same site on the RTR thus facilitating circularisation of one complete genome via the cohesive ends. Circle formation protects the T5 DNA from the degradative action of the RecBCD nuclease and eventually leads to rolling circle DNA replication.

## 1. Introduction

Bacteriophage T5 was one of the original seven “T-phages” isolated by Max Delbrück and his collaborators in the 1940s. The molecular biology of phage T5 has been reviewed (1) and other reviews specifically covered the First Step Transfer (FST) region of phage T5 (2, 3 4). T5 has a linear genome of 121,752 bp with large terminal repeats (Left Terminal Repeat, LTR and Right Terminal Repeat, RTR) each comprising 10,139 bp (5). T5 differs from most phages in that it injects its DNA in two steps; firstly, the First Step Transfer (FST) which is the left-most 7.9% and then, following pre-early gene expression, the remainder of the genome (Second Step Transfer, SST). The expression of two genes, *A1* and *A2*, is necessary for the SST DNA injection. The FST region contains the injection stop signal (*iss*) which is the locus at which DNA injection stops (3, 4, 6). The FST DNA can be separated from the SST DNA by mechanical shearing of the T5/bacterium complex, usually in the presence of cloramphenicol to prevent protein synthesis (2). It was hypothesized that *iss* could be a special nucleotide sequence on the DNA. The position of the region containing *iss* was calculated and then cloned and sequenced (6). The *iss* locus lies within the left end of a 1662 bp non-coding region which contains 97 stop codons and extends from 9194 to 10856. The *iss* region is rich in a bewildering array direct repeats, inverted repeats and palindromes, (3, 4, 5, 6) and these were shown to be conserved, and in the same order, in other T5-like phages (3, 4).

The genetic characterization of *iss* is difficult since it is an essential *cis*-acting site. The phenotype of mutants in this region would be expected to be either non-conditional lethal or wild-type; depending on whether or not an essential region was affected. Thus, in both cases, mutants could not be phenotypically identified and for 50 years there seemed no way to resolve this problem. However, recent interest in bacteriophages as therapeutic agents and food preservation agents has resulted in the isolation and nucleotide sequencing of many new T5-like phages, from different environmental sources, and capable of infecting a variety of bacterial hosts. These T5-like phages have widely diverged as visualized by the extensive deletions and substitutions and mutations within their genomes. Since *iss* is an essential *cis*-acting site, it is likely to have been preserved in evolution, even between widely divergent T5-like genomes. Previous reviews (3, 4) investigated the similarity of the sequences in the *iss* region among T5-like phages, particularly with regard to the conservation of the direct repeats, inverted repeats and palindromes that form part of the *iss* region.

The only model for First-Step Transfer (Lanni, 1968) dates back more than 52 years (2) and is unsatisfactory since it is purely descriptive and makes no predictions that can be experimentally verified. The present study presents a radically different model that can be experimentally tested. All T5-like phages conserve the *A1* and *A2* genes that are responsible for SST following the injection stop at the *iss* region. It is tempting, and reasonable, to suppose (4) that A1 and A2 proteins interact, directly or indirectly, with *iss* to relieve the injection stop and thus allow second step transfer. However, there is no evidence for this and this assumption will be questioned in the discussion section. Specifically, all that is required, for second-step injection to continue, is to separate the SST DNA from the bound membrane *iss* stop signal, permitting injection to proceed.

The author (JD) is now retired from laboratory activity and the reported new data are thus entirely *in silico.* The purpose of this article is to report these new findings and to make to suggestions and hypotheses that can be experimentally tested by others.

## 2. Materials and Methods

### 2.1 Nucleotide Sequence Analysis

Nucleotide sequence analysis was performed using classical methods such as NCBI Blast analysis, EMBOSS, Galaxy Fuzznuc, and EBI Clustal.

### 2.2 Nomenclature and the T5 genetics

Nomenclature is important in understanding the genetic map of T5 shown in Figure. 1. The left terminal repeat (LTR) of 10,139 bp is identical in nucleotide sequence to the right terminal repeat (RTR). T5 DNA is transferred in two steps: the First Step Transfer (FST) and the Second Step Transfer (SST). The FST region is contained between the left terminus of the molecule and the point of shear of the phage/bacterium complex in a blender and is estimated about 7.9% of the genome (roughly the leftmost 9618 bp). In the absence of A1 and A2 proteins, the *iss* sequence prevents SST transfer; presumably by an *iss* DNA/bacterium interaction (discussed below). The second step transfer (SST) region comprises the remainder of the unique part of the genome, as well as the RTR and a small part of the right end of the LTR (between *iss* and the right end of the LTR).

**Figure 1.**
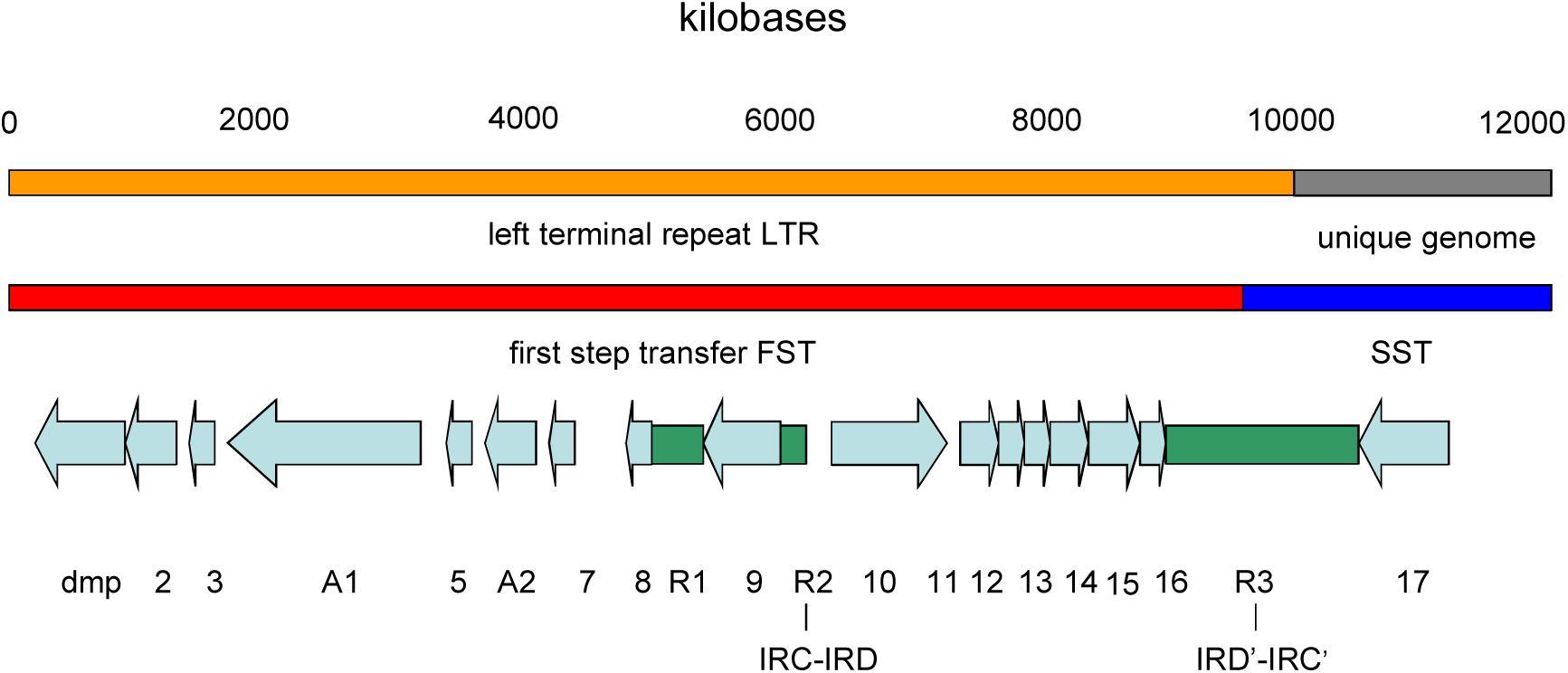
Genetic map of the LTR of phage T5. Only the left 12,000 bp of the 121,750 bp phage T5 genome are shown. The top line shows the position of the LTR (orange). The line 2 shows the FST (red) and the left end of the SST (dark blue). Line 3 shows the positions of the ORFs (pale blue) corresponding to the Paris-Orsay nucleotide sequence (AY692264.1). Line 3 also shows the positions of the repeat regions R1, R2 and R3 (green). R2 and R3 respectively contain the inverted repeats IRCspacerIRD and IRD’spacerIRC’ (IRD’spacerIRC’ is the inverted repeat of IRCspacerIRD). The injection stop signal *iss* is associated with IRD’spacerIRC’ and probably also with IRCspacerIRD. Repeat regions R1 and R2 are separated by the insertion of a HNH endonuclease gene (ORF 9). This insertion is not present in most T5-like phages.

### 2.3. T5-like phages sequences used in this study

The collection of 25 T5-like phages sequences used in this study is shown in Table 1. All of these phages carry the *dmp, A1* and *A2* genes, in this order, in the left end of the LTR. They all also contain the large untranslated region containing the *iss* locus in the right end of the LTR; discussed in detail below (4). Some true T5-like phage isolates are nearly identical, and, in these cases, only one example of each was chosen for analysis. Thus, chee24 is very similar to pork27, pork2, saus47N, saus111K, poul124; while chee130_1 is almost identical to saus132, poul149, chee158, cott162 and saus176N. Similarly, DT57C is almost identical to DT571/2 in the LTR region.

**Table 1.** Phage nucleotide sequences used in this study

| Phage Name | NCBI Accession Number | Reference |
| --- | --- | --- |
| Phage T5 (Moscow) | AY543070.1 | unpublished |
| Phage T5 (Hangzhou) | AY587007.1 | (5) |
| Phage T5 (Paris Orsay) | AY692264.1 | unpublished |
| *Enterobacteria* phage EPS7 | CP000917.1 | (7) |
| *Yersinia* phage phiR201 | HE956708.2 | unpublished |
| *Enterobacteria* phage SPC35 | HQ406778.1 | (8) |
| *Escherichia* phage AKFV33 | HQ665011.1 | (9) |
| *Escherichia* phage FFH1 | KJ190157.1 | (10) |
| *Salmonella* phage Stitch | KM236244.1 | (11) |
| *Enterobacteria* phage DT57C | KM979354.1 | (12) |
| *Salmonella* phage Shivani | KP143763.1 | (13) |
| *Escherichia* phage chee24 | MF431730.1 | (14) |
| *Escherichia* phage chee130_1 | MF431736.1 | (14) |
| *Salmonella* phage BSP22A | KY787212.1 | unpublished |
| *Escherichia* phage Gostya9 | MH203051.1 | (15) |
| *Pectobacterium* phage My1 | JX195166.1 | (16) |
| *Pectobacterium* phage DU_PP_V | MF979564.1 | unpublished |
| *Klebsiella* phage IME260 | KX845404.2 | unpublished |
| *Klebsiella* phage Sugarland | MG459987.1 | (17) |
| *Providentia* phage PR1 | KY363465.1 | (18) |
| *Salmonella* phage Sw2 | MH631454.2 | unpublished |
| *Salmonella* phage Seafire | MK050846.1 | (19) |
| *Proteus* phage Stubb | MH830339.1 | (20) |
| *Salmonella* phage Sepoy | MK728825.1 | (21) |
| *Salmonella* phage Seabear | MK728824.1 | (22) |
| *Serratia* phage Slocum | MN095770.1 | (23) |
| *Escherichia* phage BF23 | Mei Liu, Personal communication. | unpublished |
| Phage T5 injection stop signal | M16226.1 | (6) |

Genbank contains many more sequences that have homology to T5 but only those in Table 1 were retained as being true T5-like phages. Others seem to be the product of sequencing errors and/or sequence assembly errors; and have (e.g.) the *dmp, A1*, and *A2* genes in the late region of the genome or have late genes in the LTR. This seems due to the fact that sequence assembly software has difficulty in processing the presence of the large terminal repeats.

### 2.4. The nucleotide sequence of the first step transfer region

There are differences between the 3 published T5 nucleotide sequences (Hangzhou AY587007.1 Paris-Orsay AY692264.1, and Moscow AY543070.1) and these were analyzed in detail by Davison (3). These differences make it difficult to interpret the data for the identification and nomenclature of the ORFs in the FST region. In particular, the Hangzhou sequence is missing 75 bp, starting at coordinate 5599, compared to Paris-Orsay and Moscow sequences. This, and other smaller differences, results in a lack of a complete correlation of the Hangzhou ORFs compared to the Paris-Orsay and Moscow sequences (which are very similar, though not identical).

Consequently, the Paris-Orsay AY692264.1 sequence coordinates have been used for ORF analysis. However, the Hangzhou (AY587007.1) nucleotide sequence publication contains an invaluable detailed analysis of the regions containing repeats, inverted repeats and palindromes and these coordinates are used for discussion of the repeat regions.

## 3. Results

### 3.1. Phylogenetic relationships of T5-like phages

A phylogeny analysis (Figure 2) using the Dmp protein shows that most T5-like phages form a single clade (Type1) which includes T5, BF23, FFH1, AKFV33, chee24, Shivani, Gostya9, DT57C, EPS7, BSP22A, chee130-1, Stitch, phiR201, SW2, Seabear and Seafire. Type 2 phages (My1, and DU_PP-V) and Type 3 phages (IME260 and Sugarland) are on separate branches that are evolutionarily distant from each other and from Type 1 phages. Another phage Slocum is closest IME260 and Sugarland and has been classified as Type 3 with these phages. A fourth clade (Type 4) has 2 members PR1 and Stubb and is very distant from the other Types.

**Figure 2.**
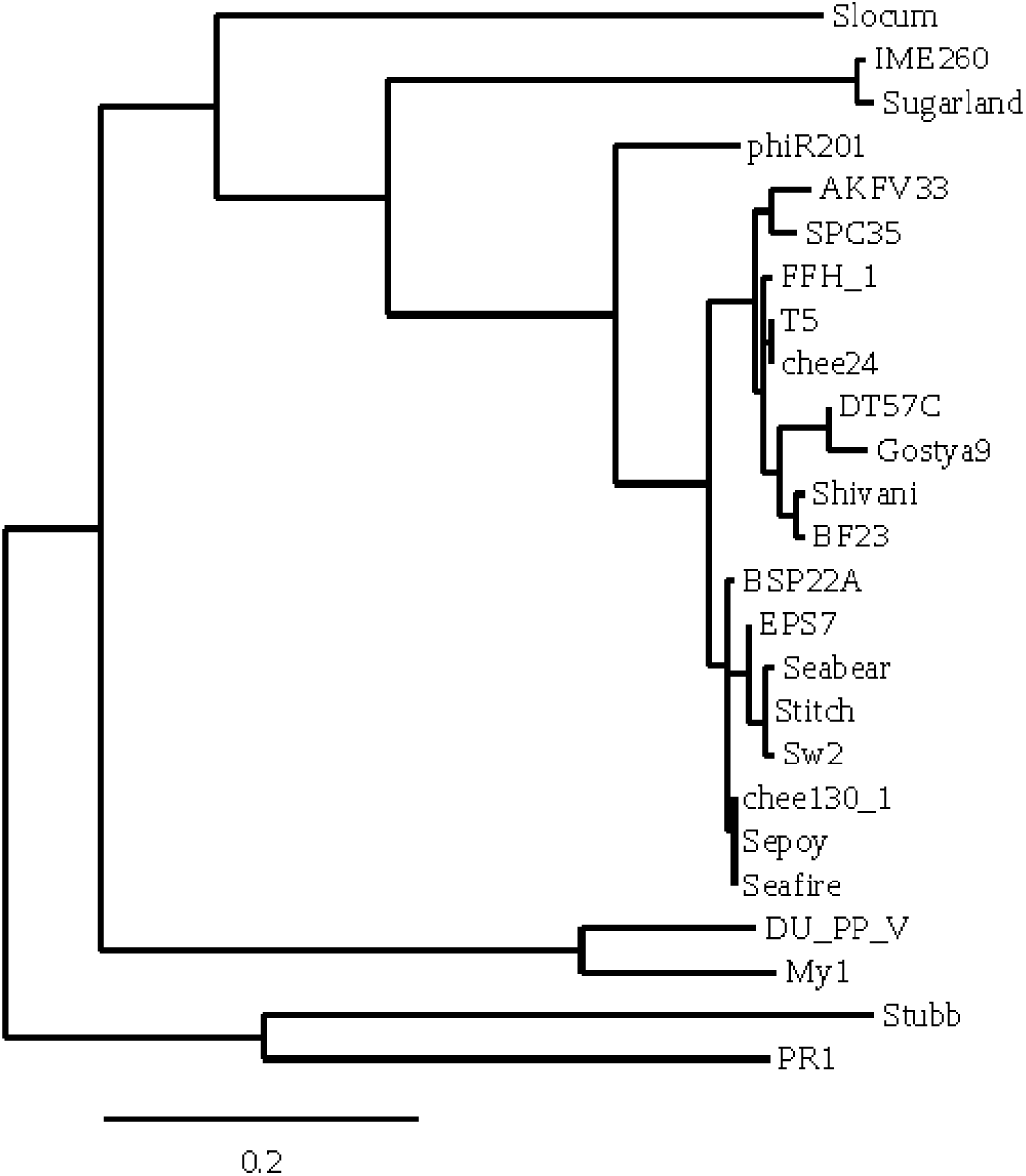
Phylogenetic relationships of the T5-like phages. The relationships were determined with the deoxynucleoside 5’-monophosphatase amino acid sequences of the corresponding phages, using the Phylogeny.fr program.

Members of the Type 1 group have many small differences between themselves, yet the difference between Type 1 and Types 2 and 3 is more profound since in Types 2 and 3 the entire region corresponding to the T5 region ORFp01 1 to ORFp015 is missing and is replaced by other nucleotide sequences containing ORFs that are read in the opposite direction to those of T5 (Fig.1).

The Type 4 phage, PR1, is very different from the others and Oliveira et al (18) suggest that PR1 forms to a new species within the *Siphoviridae* family. The newly isolated phage Stubb is tentatively assigned to Type 4 and this is supported by the pattern of its *iss* region (discussed below).

A word of caution is, however, necessary in comparing these various largely uncharacterized T5-like phages to study the injection stop signal *iss*. Only two phages, T5 and BF23, that have been studied genetically and biochemically and two-step injection has been verified only in these two phages. It is thus an assumption, based on the presence of the *A1* and *A2* genes and on the common structure of the *iss*, regions that all T5-like phages use two-step injection.

### 3.2. The ORFs in the T5 FST region

It was interesting to compare the ORFs in the FST region since this would give an indication of which FST ORFs are essential for phage growth. It was further important to establish the similarities and differences between these phages for comparison with the similarities and differences found in the non-translated *iss* region. As will be noted in the discussion, the non-translated *iss* region is more rigorously maintained than are many of the coding regions.

#### 3.2.1. The relationships between the ORFs of T5-like phages

The T5 coding region 1 to 6022, corresponding to the rightward transcribed genes (Figure 1), is generally conserved, with several variations, in most T5-like phages. The presence, or absence, of these ORFs is shown in the Supplemental Files.

In contrast, the coding region 6209 to 9194 between Repeat Region 2 and Repeat Region 3 of T5 shows highly divergent nucleotide sequences, with many deletions/insertions, between the different T5-like phages. In this region, phage T5 has almost no DNA homology to the Type 2 (My1 and DU_PP_V) and Type 3 T5-like phages (IME260, and Sugarland). Furthermore, it is transcribed from the opposite strand.

Phage DT57C is instructive since it has the smallest LTR (8081 bp) containing only 11 ORFs (compared to only 16 in T5 and 22 in phage My1) due to a deletion (compared to T5) that removes T5p008, T5p009, T5p010, and T5p011. Gostya9 similarly has a large deletion in this region.

Phage My1 (Type 2) is also interesting since it has the longest LTR (12854 bp) containing 22 ORFs. Many of these ORFs are unique to My1, or are shared only with the other Type 2 phage DU_PP_V. Both My1 and DU_PP_V lack T5 ORFs T5p011 through T5p015 though they retain T5p016. ORF My1_018 is curious, being homologous to ORFs from DU_PP_V, but also to some Type 1 phages (chee24, Shivani, SW2, EPS7, and Stitch); but having no homolog in T5.

The original mutant hunts (2) identified only 2 genes (*A1* and *A2*) essential for phage viability in the LTR. Another gene *dmp* (encoding a deoxynucleoside 5’-monophosphatase), is not essential for growth. These genes are shown in Figure 1, together with the other putative ORFs in the T5 FST region. It is not excluded that other essential genes could have been missed in early mutant searches, particularly since the searches were not extensive and since some FST ORFs are quite small. In particular, the T5 FST region seems to have more functions than identified genes to accomplish them: injection stop, restart of phage injection, degradation of host DNA (but not T5 DNA), shut-off of host RNA and protein synthesis, shut-off of pre-early genes, protection against restriction enzymes and CRISPR and protection against RecBCD nuclease. Thus, the FST region may contain unidentified genes that may facilitate these functions. For this reason, it was interesting to compare the ORFs of different T5-like phages since important ORFs are likely to be conserved, while non-essential ORFs could be lost. Previous studies (3, 4) found that only genes *dmp, A1, A2* and T5p007 were common to all T5-like phage nucleotide sequences. Gene T5p007 has no known function but could be a candidate for the unknown gene conferring restriction insensitivity (3, 4, 24, 25, 26). A more detailed summary of the ORF content of the pre-early region of the T5-like phages is given in the Supplemental Files. The *A1* and *A2* genes are concerned with two-step DNA injection and are thus the only known genes relevant to this publication.

#### 3.2.2. The *A1* gene

*A1* mutants do not inject SST DNA, cause degradation of host DNA, or shut-off of host RNA and protein synthesis or shut-off of T5 pre-early gene activity. The A1 and A2 proteins form a heterodimer, which is consistent with a role of both of these proteins in SST injection. A1 protein is associated with membranes where it accounts for up to 10% of the newly synthesized membrane proteins following infection (27). Membrane association is consistent with a role in SST injection since, as shown below, it is likely that the FST DNA is membrane bound.

*A1* mutants do not degrade host DNA, suggesting perhaps a nuclease activity. However, apart from one preliminary report (28), no nuclease activity has been reported for the A1 protein or any other pre-early protein. One possibility, discussed in detail below, is that *A1* codes for a site-specific restriction endonuclease. It seems unlikely that a search for a site specific restriction enzyme has been conducted since this would necessitate a specific DNA substrate carrying the appropriate restriction sites, special analysis by gel electrophoresis and perhaps adding ATP and *5*-adenosylmethionine to the reaction mixture. The restriction enzyme hypothesis for the A1 protein, suggests that restriction sites are present in the host DNA but not in the FST DNA, would explain the age-old enigma of how T5 degrades host DNA without degrading its own DNA: this would be simply due to the restriction sites carried by those DNA molecules.

#### 3.2.3. *A2* gene

Like *A1* mutants, *A2* mutants do not inject SST DNA. A2 protein has been purified and shown to be a dimer that binds double stranded DNA (29, 30). It also associates with A1 protein, which is a membrane binding protein..

Cloning of the *A2* gene in phage λ shows it has no deleterious effect on λ replication and the 1-T5 *A2^+^* hybrid is able to complement a T5 *A2* mutant (31). Like phage T5, a λ phage carrying the *A2^+^* gene is inhibited in *E. coli* carrying the *ColIb* plasmid. This shows that only the A2 protein is responsible for this abortive infection (32). Little is known of the mechanism of inhibition by ColIb.

### 3.3. The structure of the untranslated FST Repeat Regions

The LTR of T5 carries 3 untranslated repeat regions:

1. Repeat Region 1, (629 bp) 4682 to 5310
2. Repeat Region 2, (188 bp) 6022 to 6209
3. Repeat Region 3, (1.2 kb) 9194 to 10409

These 3 repeat regions are highly conserved with the same order of repeats, inverted repeats and palindromes between almost all of the T5-like phages indicating that they serve some important functions (3, 4). It is instructive to contrast the high level of conservation of the untranslated repeat regions 2 and 3, among the T5-like phages, to the much lower level conservation of many LTR ORFs; where only *dmp, A1, A2* and ORFp007 are consistently conserved.

Repeat Region 1 is present in almost all T5-like phages and contains IRA and IRA’ and IRB. However, its locus is too near to the left end of the LTR to be considered as a candidate for *iss* which should logically be in Repeat Region 3.

Repeat Region 2 contains the IRC, IRD and IRB’. Repeat region 2 is also not a prime candidate for the *iss* signal. On the other hand, it contains inverted repeats IRCspacerIRD (6022 to 6152) which are nearly identical to IRD’spacerIRC’ in Repeat Region 3 and this latter is a probable candidate for *iss*. In the Discussion section, it will be argued that both IRCspacerIRD and IRD’spacerIRC’ are part of the *iss*, despite the large distance between them (this distance being highly variable between different T5-like phages).

Repeat Regions 1 and 2 are separated in phage T5, Shivani and FFH1 by the insertion of the HNH endonuclease gene (ORF 9 in Fig.1). Most T5-like phages do not carry this insert and it is obvious that Repeat Regions 1 and 2 were originally a single repeat region in the ancestor of T5-like phages.

#### 3.3.1. Repeat Region 3: Repeats, Inverted Repeats and Palindromes

Repeat Region 3, (1.2 kb) 9194 to 10409 is mostly in the LTR but overruns into the unique genome by 270 bp. Repeat Region 3 is extremely complicated, rather like a Matryoshka doll, or Russian nesting doll, where one structure hides another, which in turn hides another and then another (Figure 3). It contains palindrome palC, palindrome palD, inverted repeat IRD’, direct repeat DRE1, direct repeat DRB copy 1, direct repeat DRB copy 2, inverted repeat IRC’, direct repeat DRE2, direct repeat C copy 1, palindrome palE, direct repeat DRC copy 2, direct repeat DRC copy 3, palindrome palH, palindrome palG, direct repeat DRC copy 4, palindrome palI, palindrome palJ, 9 copies of direct repeat DRD, and direct repeat DRA copy 2, and copy 3.

**Figure 3.**
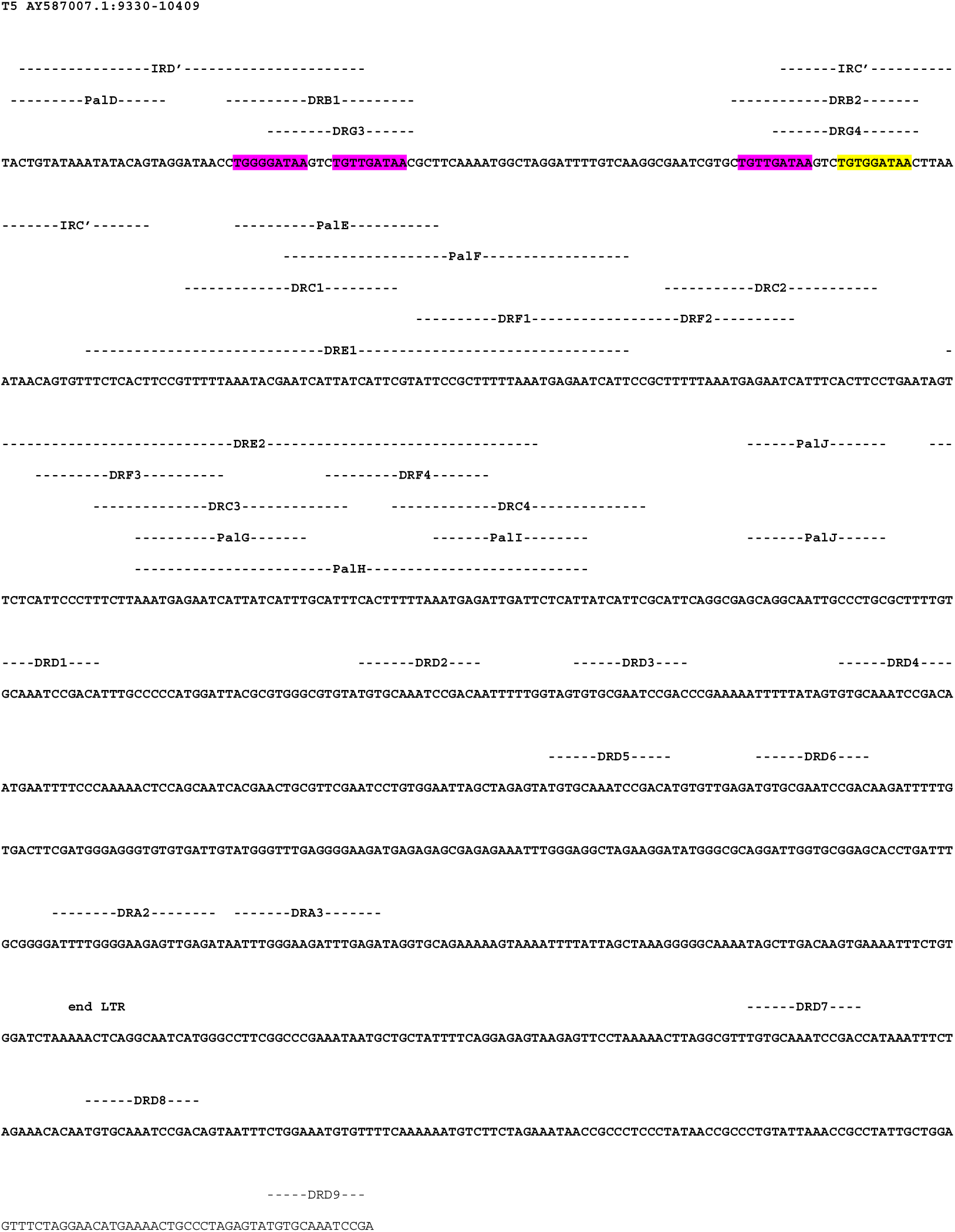
The repeats, inverted repeats and palindromes in Repeat Region 3 of T5. The nucleotide sequence of the untranslated region (from coordinates 9330 to 10409 of AY587007.1) is shown together with the direct repeats (DR), inverted repeats (IR) and palindromes (Pal) it contains. The 131 inverted repeat IRD’spacerIRC’ (9332 to 9462) comprises IRD’, IRC’ and the region between them. The other partner of this inverted repeat is the 131 bp IRCspacerIRD (coordinates 6022 to 6152), not present on this Figure, but is shown on Figures 1 and 4). The approximately calculated shear-point for the FST would be at coordinate 9618 in DRE2. The positions of the DnaA (yellow) and DnaA* (magenta) protein binding sites are shown (see also Figure 4). Because of the strand orientation, the DnaA protein binding sites are shown as the reverse complement. The end of the LTR is indicated.

Repeat Region 3 is in the correct position, just to the left of the shear site, to contain the *iss* site and is shown in Figure 3. It comprises at least 3 major kinds of repeat sequences that have been characterized previously (3, 4):1) IRD’spacerIRC’ (9332 to 9462, 131 bp), 2) two DRE repeats (66 bp), 3) a series of nine DRD repeats.

The use of the word ‘spacer’ in the naming of these repeats is an attempt to maintain the useful IRD’ and IRC’ nomenclature of Wang et al (5). It should not be interpreted to believe that the ‘spacer’ is without function. Indeed, evidence is given below that the ‘spacer’ and the inverted repeats contain functionally important elements.

#### 3.3.2. Repeats Inverted Repeats and Palindromes in Repeat Region 3

It was shown (6) that the Repeat Region 3 contains a multitude of repeats, inverted repeats and palindromes and this was confirmed by sequencing of the entire genome (5). However, during the course of this investigation some new repeats and inverted repeats were discovered. These were documented in a previous publication (4) and can be visualized in Figure 4. They are described in the Supplemental Files

**Figure 4.**
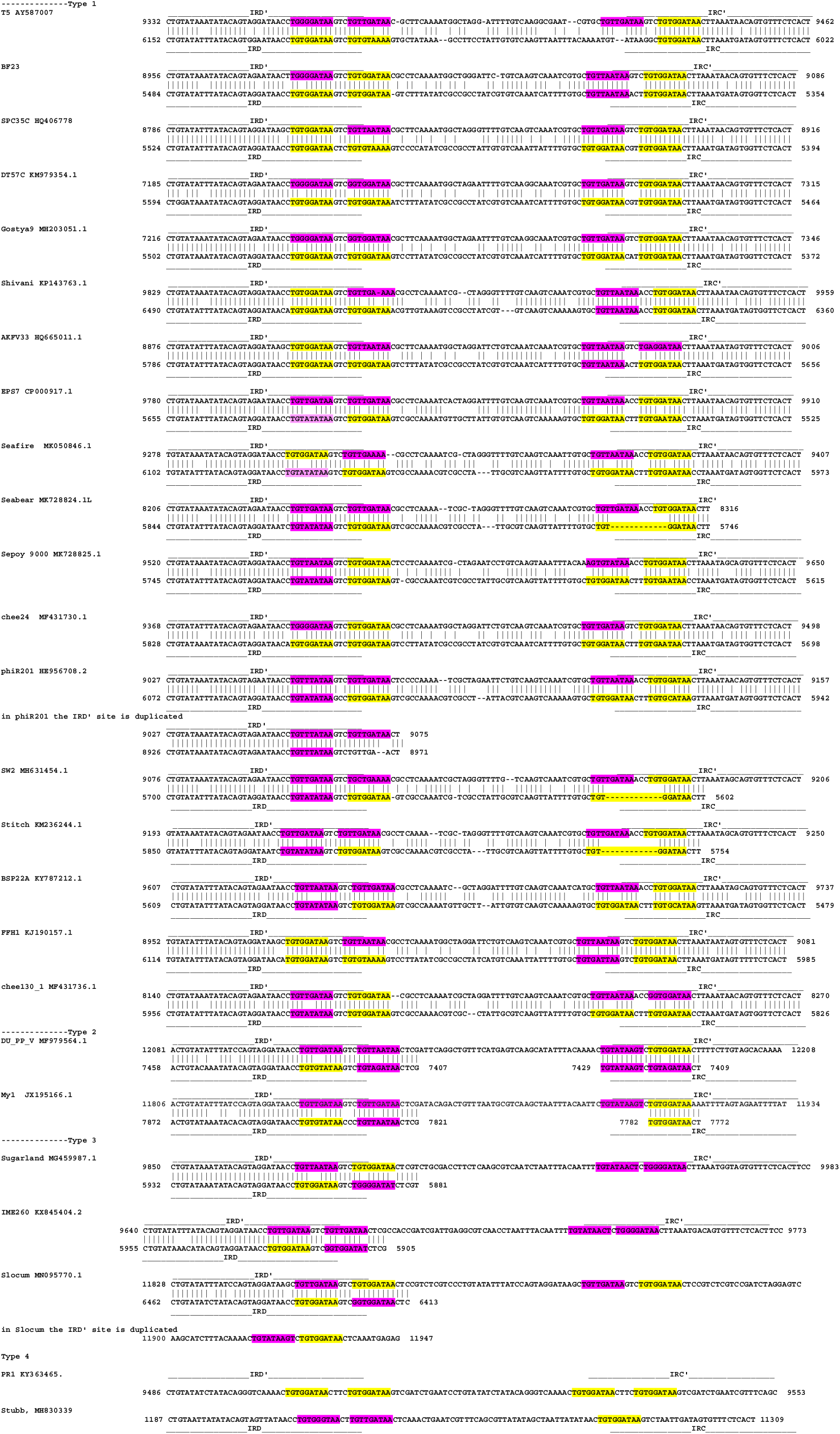
DnaA protein binding sites in the IRD’spacerIRC’ and IRCspacerIRD inverted repeats. The top line of this figure corresponds to the top line of Figure 3. The inverted repeats IRD’spacerIRC’ (top line of each phage) is drawn in the classical direction, 5’ to 3’; from left to right. The inverted repeat IRCspacerIRD inverted repeats (bottom line of each phage) is drawn as the reverse complement 3’ to 5’. Figure 4 is drawn in this way only to illustrate the homology and does not imply that these sequences exist in this way *in vivo.* Since these are inverted repeats the two sequences are thus largely homologous. The phage sequences are derived from those in Table 1 and the coordinates of each inverted repeat are shown at the beginnings and ends of the lines. Gaps in the sequences indicate non-homology due to missing nucleotides. The DnaA and DNA* bindings sites are shown. Consensus DnaA binding sites TT(A)/(T)TNCACA (TTATCCACA, TTATTCACA; TTTTACACA, TTATGCACA) are shown in yellow and non-consensus DnaA binding sites (for example TTATCAACA and TTATTAACA) are shown in magenta. It should be noted that, because of the strand orientation, the DnaA protein binding sites and the DnaA* protein binding sites are drawn as the reverse complement; so that, for example, TGTGGATAA is equivalent to TTATCCACA but on the complementary strand.

#### 3.3.3. DnaA boxes

It was shown in 1987 (6), that the IRD’spacerIRC’ region contains one consensus DnaA box TTATCCACA, and also contains 3 variant DnaA boxes; TTATCAACA (present twice) and TTATCCCCA (present once) (Figure 3). Consensus DnaA boxes are defined according to of Schaper and Messer (33) TT(A)/(T)TNCACA). For convenience the variant sites, which differ from the consensus by one or two nucleotides, will be collectively referred to below as DnaA* boxes.

This observation prompted a search for DnaA boxes and DnaA* boxes in the equivalent positions in other T5-like phages. The results are shown in Figure 4. It can be seen that consensus DnaA boxes and DnaA* boxes, are present in the IRD’spacerIRC’ region of all T5-like phages.

Figure 4 also shows the IRCspacerIRD and it can be seen that DnaA boxes and DnaA* boxes are also common in the IRCspacerIRD region. This Figure is drawn as the reverse complement of the IRD’spacerIRC’ region (since these are inverted repeats) and it should be noted that this is done purely for representational purposes and is not meant to imply that such a structure exists in reality.

It is interesting that the DnaA and DnaA* boxes, for Types 1, 2, and 3, T5-like phages) are always in pairs separated by 3 bp (which is reminiscent of the clustering of DnaA sites around the oriC region). They always in the same position relative to the two IRD’spacerIRC’ and IRCspacerIRD inverted repeats with one half in the inverted repeat and the other half of other in the spacer (Figure 4). Indeed, almost all T5-like phages carry eight DnaA boxes or DnaA*boxes in the FST region: with 2 in IRD’, 2 in IRC’, 2 in IRC and 2 in IRD. T5 itself is an exception having only 7 sites (the 8^th^seems to have been destroyed by a 2 bp deletion in IRC). About half of the DnaA boxes shown in Fig 4 are consensus DnaA boxes (TTATCCACA (the most common);

TTATTCACA, TTTTACACA; TTATGCACA) and the other half are DnaA* boxes (the most common being TTATCAACA and TTATTAACA) which differ from the consensus DnaA boxes by only 1 or 2 bp. It should be noted that consensus site (30) was based upon the binding of naturally occurring DnaA boxes around *oriC, rpoH* and *mioC.* Thus, it is not known whether the DnaA* boxes would bind DnaA protein more weakly or more strongly. In addition, DnaA protein binding properties concerns not only the nucleotide sequence but also ATP, the context of the neighboring nucleotides, the interaction with bending proteins IHF and FIS and cooperativity between DnaA proteins.

The spacer distance between IRD’ and IRC’ and between IRC and IRD is always 52 bp for all T5-like phages (see below). The significance of this is unknown, but the double helix makes one complete turn about its axis every 10.4-10.5 base pairs. Thus, they are separated by 5 (52/10.4) helix turns which may ensure that certain features are always on the same side of the DNA (for example for protein binding). The spacer region is also a region of DNA bending which again may indicate protein binding

The presence of the 3 deletions in IRC of Type 1 phages (T5, Stitch and SW2) seems to indicate that not all eight DnaA are absolutely essential. This is supported by the observation that Type 2 and 3 lack part of IRC. It is noteworthy that the missing DnaA boxes are always in the IRC region; as if this region only has secondary importance to the *iss* function. Type 4 T5-like phages carry only IRD’spacerIRC’ and are completely lacking in IRD and IRC.

Finally it should be noted that two phages phiR201 and Slocum have duplications of the IRD’ region.

DNA replication origins have been shown to be near the centre of the T5 DNA molecule (34, 35, 36). It thus seems unlikely that the DnaA boxes in the FST DNA are concerned with DNA replication. Furthermore, T5 replication functions are carried by the SST region and thus are not expressed during the FST period.

### 3.4. Rolling circle formation

Circle formation in T5 is not well understood but has been presumed to be initiated by some kind of recombination between LTR and the RTR. Irrespective of the mechanism a recombination event would necessarily involve the loss of one complete terminal repeat and indeed the circles observed by electron microscopy correspond to the whole T5 DNA molecule minus one terminal repeat (31, 32, 33). As part of the model described below, it is suggested below that this recombination may be facilitated by cohesive ends generated by cleavage of the A1/A2 restriction enzyme and may be part of the SST injection process.

## 4. Discussion

### 4.1 Events following T5 infection

#### 4.1.1 Phage injection and injection stop

Phage injection takes place from the left hand end of the molecule (37) and is initiated by the simple interaction of the bacteriophage with the host outer-membrane receptor protein FhuA (previously called TonA) which is concerned with ferrichrome transport and also acts as a receptor for phages T1 and D80 (38). *In vivo*, but in the absence of protein synthesis, DNA transfer stops when only 7.9% has been injected. However FhuA seems not to be responsible for the injection stop since complete genome injection can be demonstrated *in vitro* using proteoliposomes containing FhuA protein (39). This is further supported by the observation that a different T5-like phage, BF23, which also undergoes two-step injection, uses a different receptor; BtuB; the *E. coli* vitamin B12 outer membrane transport protein (40). Phages DT57C and EPS7 also uses the BtuB receptor (6, 41). Other T5-like phages described in this article infect very different bacterial hosts and possibly also use different receptors.

Thus the question as to the host protein responsible for the injection stop is unsolved. However, the presumptive *iss* regions of T5, and all of its relatives, carry DnaA boxes to which DnaA protein binds (Figure 4). DnaA protein is ubiquitous among almost all bacteria and is found attached to the plasma membrane and forms helical structures along the longitudinal axis of the cell (42). The DnaA protein has a 35-fold higher density in the plasma membrane than in the cytosol. As will be discussed below, it seems possible that FST injection stops because the membrane bound DnaA protein binds to the DnaA boxes of the *iss* region.

#### 4.1.2 DNA degradation

Within the first few minutes of T5 infection the host DNA is degraded to nucleotides and then to free bases which are excreted from the cell. Host DNA degradation also occurs in *A2* mutants, but not in *A1* mutants; suggesting that the A1 protein is involved in host DNA degradation. However, no nuclease activity has been reported to be associated with the A1 protein. The rapid degradation of host DNA also raises the question of how T5 DNA itself escapes this degradation; particularly since T5 DNA has no special characteristics (special bases or glycosylation) that distinguish it from that of the host. One possibility (discussed below) is that the *A1* gene encodes a restriction enzyme active on sites in the *E. coli* genome; while the T5 FST DNA carries no such sites. According to this hypothesis the restriction fragments from the *E. coli* genome would eventually be solubilized by the RecBCD nuclease (next section).

#### 4.1.3 Protection against RecBCD exonuclease

To quote Dillingham and Kowalczykowski (43) “*any phage that exposes free DNA ends as part of its life cycle must find a means to evade destruction by RecBCD”.* Unlike the restriction insensitivity function (see below), protection against RecBCD nuclease cannot be explained by newly synthesized proteins coded by the FST region. The left end of T5 DNA enters the cell first and should immediately be exposed to RecBCD nuclease before any FST proteins could be synthesized. Many phages such as Mu, P22 and T4 have a terminal protection protein attached to the ends of their DNA in the phage particle and this is injected into the host along with the DNA. It was suggested previously (3, 4) that T5 may also carry a similar protection protein but this has not yet been investigated by testing the infectivity of T5 DNA extracted without the use of denaturing agents such as phenol. Whether T5 encodes a RecBCD inhibitor (as do many phages) is unknown.

#### 4.1.4 Turn-off of host RNA and protein and FST RNA and protein synthesis

Both host and phage protein and RNA synthesis are turned off after about 5 minutes. The A1 protein (but not the A2 protein) is necessary for this turn-off. It is probable that turn-off is a consequence of host DNA degradation.

#### 4.1.5 Restriction Insensitivity

Restriction insensitivity was discussed at length in a previous reports (3, 4, 24, 25, 26) and will not be repeated here since it is outside of the scope of this article. However, recent publication (44) investigated the effect of the CRISPR-cas system on T5. It was found that T5 was insensitive to *E. coli* CRISPR-cas strains targeting the early and late regions of T5. However, T5 was sensitive to *E. coli* CRISPR-cas strains targeting the FST region. Though the authors were unaware of the earlier *EcoRI* restriction insensitivity papers, this CRISPR situation exactly parallels the results with restriction by *EcoRI* restriction enzyme; whereby *EcoRI* sites in the early and late regions are insensitive to *EcoRI* restriction and *EcoRI* sites in the pre-early FST region were cleaved by *EcoRI* (41, 42, 43). This suggests that CRISPR-cas may, perhaps, be added to the list of endonucleases that are protected by a T5 pre-early function.

### 4.2 Repeat region 3 and the injection stop signal

The FST region of T5 is defined by the site of shear of T5 infected cells in a blender. The usual estimate of the FST region is left-most 7.9% (roughly 9618 bp from the left); which falls within DRD2 repeat (Figure 3) and iss is thought to be to the left of this shear site (6). Inspection of this region reveals the large inverted repeat IRD’spacerIRC’ (131 bp) and the immediately adjacent 66 bp DRE1 and DRE2 repeats, as shown in Figure 3. These are, individually, or together, likely candidates for iss and are present in all T5-like phages. This high degree of conservation is all the more remarkable given the almost complete lack of conservation of the ~3 kb coding region immediately to the left of Repeat Region 3 (see above and Fig. 1). The IRD’spacerIRC’-DRE cluster is extremely complex containing within it many repeats and palindromes shown in Figure 3. It is impossible to guess the function(s) of these, though their extreme conservation, among all T5-like phages, suggests their importance (3, 4).

### 4.3 Repeat region 2 and the injection stop signal

Repeat region 2 begins at 6022 and ends at 6209 and thus is too far leftwards from the shear site 9618 to be considered as *iss.* Yet the 131 bp inverted repeat IRD’spacerIRC’ of Repeat Region 3 is also present, (in inverted order, as IRCspacerIRD), in Repeat Region 2. This cannot be a coincidence since IRCspacerIRD is present at similar positions in almost all T5-like phages. This, almost inevitably, leads to the conclusion that IRCspacerIRD and IRD’spacerIRC’ are part of a functional complex and that they must both be part of the *iss* injection stop process. This hypothesis is reinforced by the finding of common elements between IRC and IRD and IRD’and IRC’; namely the DRB and DRG repeats and DnaA boxes discussed below).

One observation may elucidate this situation: while investigating the FST injection of T5, Rhoades (45) observed two FST bands on the gels. The first and largest corresponded to the classical FST band of about 9618 bp, but a second band was present at about 5846 bp which (given the possible error measurements in 1979) is very close to the position of IRC at 6022. The authors did not have an explanation for this band. However, this raises the intriguing possibility that the FST DNA may attach to the membrane in two places; at IRD’spacerIRC’and at IRCspacerIRD (coordinates ~9618 and ~6022.)

### 4.4 The Classical model of T5 Injection

The classical Lanni model (2) for the two-step injection of T5 states that injection stops (for unknown reasons) and then resumes (for unknown reasons) under the action of the newly synthesized *A1* and *A2* products. While this model has stood the test of time (52 years!), and no other model has been proposed, it is unsatisfactory since it simply describes the phenomenon.

The classical model makes only one prediction, namely that 100% of the intact genome enters the host cell. Two studies investigated the replication of T5 DNA, using electron microscopy, and both observed circular DNA molecules carrying replication forks and the length of the circle was that of the T5 DNA minus the length of one terminal repeat. This confirms the idea that circular molecules are formed by some kind of recombination of the LTR and RTR resulting in the elimination of one repeat. These circles are probably rolling circle precursors.

However, whereas Bourguignon et al. (31) observed full length linear T5 DNA carrying replication loops, while Everett (33) also observed linear molecules carrying replication loops but these were never full length, being 12% shorter than T5 DNA and thus could not be replication intermediates. In view of the fundamental disagreement between these two results, the question of whether intact full length T5 DNA enters the cell remains unresolved (4).

One major problem with the Lanni model for two-step injection is that it provides no hypothesis as to why the DNA should stop. Another major problem is that it makes no attempt to reconcile the facts that the A1 protein (directly or indirectly) degrades the host DNA yet spares T5 FST DNA. The A1 protein, together with A2 protein, is also required for SST injection but there is no hypothesis as to how this is achieved. These ambiguities must be resolved and a new hypothesis for T5 injection is thus needed.

### 4.5 A new model for T5 injection stop

The present study reveals the vast complexity of repeat region 3 which contains within itself a multitude of overlapping repeats, inverted repeats and palindromes. The fact that the untranslated *iss* region is highly conserved, between different phage genomes that are not themselves well conserved in their coding regions, attests to its importance. Its locus, just to the left of the shear site, suggests it is likely to be (wholly or partly) the *iss* region. The conservation of this region suggests that cannot it be greatly altered without destruction of the *iss* function. Furthermore the present study demonstrates the presence of 8 precisely organized putative DnaA boxes; one pair of DnaA boxes being associated with each of the IRC, IRD, IRD’ and IRC’ inverted repeats. Each FST region of all T5-like phages has eight DnaA boxes (with rare exceptions; including T5 itself, which has only 7, Figure 4) that are arranged in pairs and separated by 3 nucleotides; which is reminiscent of the precise clustering of DnaA sites around the *oriC* region of *E. coli.*

It has been shown that, while DnaA protein is purified from the cytosol as a soluble protein, the vast majority is bound to the plasma membrane, with a 35-fold higher density in close proximity to the cell membrane than in the cytosol (46). It is unknown whether these membrane bound DnaA proteins would interact with the DnaA boxes of T5 which must pass within close proximity during the injection process. This could perhaps be tested using a temperature sensitive *dnaA* mutant at the non-permissive temperature or by using a *dnaA* deletion host.

The presence of 8 DnaA boxes in the IRCspacerIRD and the IRD’spacerIRC’ regions of the FST DNA, together with the numerous repeats, and palindromes, could cause considerable bending of the DNA when bound to DnaA protein (and perhaps other proteins such a IHF and FIS). It could even form a knot, together with the IRCspacerIRD regions which could physically prevent transfer.

Given this hypothesis as to why FST injection stops, it is now necessary to know how SST injection restarts under the influence of the A1 and A2 proteins.

### 4.6 The A1 protein-restriction enzyme hypothesis

The classical Lanni model (2) of T5 two-step injection ignores the inconvenient truth, which was already know at that time, that the A1 protein is probably an endonuclease since it causes degradation of the host DNA. T5 FST DNA is not degraded, indicating that it is protected from the nuclease that destroys host DNA. The most satisfying explanation is that the A1 protein attacks sites on the host DNA that are not present on the FST DNA. This is a perfect description of a restriction enzyme that would cleave restriction sites on the host but not the FST DNA which lacks these sites.

The hypothesis that A1 protein is a restriction endonuclease explains host degradation but does not explain how it detaches the *iss* signal from the membrane bound DnaA proteins to which it is attached. It is difficult to imagine a reasonable mechanism whereby a restriction enzyme could do this; leaving the mystery of how then SST injection restarts. The present model suggests the A1 restriction enzyme, perhaps in cooperation with the *A2* gene protein, also cleaves the FST DNA at a specific restriction site to the right of the site of blockage of the membrane bound FST DNA. By cleaving the FST in this way, injection is free to continue, leaving the FST DNA behind still attached to the DnaA protein in the membrane.

Restriction enzymes often cleave palindromic sequences and it should be noted (Figure 2) that there are several palindromes in the appropriate region of the T5 LTR region (PalD, PalE, PalF, PalG, PalH, Pall and PalJ). It should also be remembered that these palindromes are also present in the RTR and thus cleavage of the LTR restriction site palindrome should also imply cleavage of the RTR restriction site when it enters the cell on the SST DNA. Restriction enzymes often cleave palindromic sequences to give cohesive ends and the presence of homologous cohesive ends in the LTR and the RTR, following cleavage, could facilitate circle formation of one complete genome. This circle formation would be a pre-cursor for rolling circle replication. This process is similar to that of phage 7 with the exception that 7 cohesive ends are already available for circle formation.

It is counterintuitive to imagine that a phage would simply cleave and abandon about 7.5 % of its genome, yet it should be remembered that this would happen anyway. At the time of circle formation it has always been considered that the LTR and the RTR fuse by some kind of recombination with the result that the equivalent of one terminal repeat is lost (31, 33). Thus the only difference is one of timing: according to this hypothesis one terminal repeat is lost earlier than previously imagined.

### 4.7 The role of A2 protein

A2 protein is also needed, together with A1 protein, for SST transfer. A2 polypeptide has been purified and shown to be a DNA binding dimer (25). The A1 and A2 proteins are known to bind together and to be membrane associated. A2 protein does not play a role in host DNA degradation. The function of the A2 protein in second step transfer is thus unknown and any suggestion would be pure speculation.

The model given above could function equally well without the participation of the A2 protein. Yet, since the A2 protein is essential for second-step injection, a role for the A2 protein participation must be suggested. One possibility, is that A2 protein could modify the A1 restriction enzyme, to which it is bound, to change its specificity to be able to recognize the restriction site located in the FST DNA. According to this hypothesis the restriction site recognized by the A1/A2 complex would be more specific than the sites on the host DNA recognized by A1 alone. A possible extension of this hypothesis would be that the A1/A2 complex (or A2 alone) could have an active role in circle formation by joining the cohesive ends of restriction cleavage sites on the LTR and the incoming RTR. This would be site-specific recombination. Such a function could be analogous to the *int* protein of phage 7 which catalyses 7 DNA integration to form a prophage; yet, when combined with *xis* protein, *int* specificity changes and causes prophage excision.

### 4.8 The role of IRCspacerIRD

The above model has considered that the *iss* signal is near and to the left of the shear site and thus IRD’spacerIRC’ is considered a likely candidate for *iss*. While this is probably correct, it does not exclude a role of IRCspacerIRD. In T5 IRDC’spacerIRD’ is 3180 bp to the right of the IRCspacerIRC, but in other T5-like phages there is great variation in this distance (for DST57C it is only 1851bp, while for BSP22A it is 4258 bp); as if the actual distance was of little importance (4). IRD’spacerIRC’ and IRCspacerIRD could both have a role as *iss*, whereby the DnaA boxes they contain could bind the FST DNA to the membrane to form a Gordian knot (perhaps including the repeats and palindromes they contain) that prevents injection. The only solution, to be rid of the knot and for injection to continue, is to cut the DNA.

### 4.9 The new restriction enzyme/DnaA protein model

The new model is summarized in Figure 5 and shows FST DnaA boxes of the *iss* region attached to the membrane bound DnaA protein (Figure 1a). The *A1* coded restriction endonuclease cleaves the host DNA thereby causing host DNA degradation. The host DNA degradation is completed by the action of the RecBCD nuclease. This degradation would shut-off of host messenger RNA and protein synthesis. The A1/A2 heterodimer also cleaves restriction sites upstream (rightwards) of the *iss* signal, permitting the transfer of the SST DNA; by removing the stop caused by the attachment of the DnaA boxes to membrane-bound DnaA protein (Figure 1b). The A1/A2 heterodimer also cleaves the corresponding site on the incoming RTR DNA and facilitates circle formation either by virtue of the cohesive ends of the restriction cleaved ends or by site-specific recombination. DNA replication leads to rolling circle formation and hence to concatenate formation. Following late gene expression and head and tail formation, the concatenate is then processed and packaged into phage particles.

**Figure 5.**
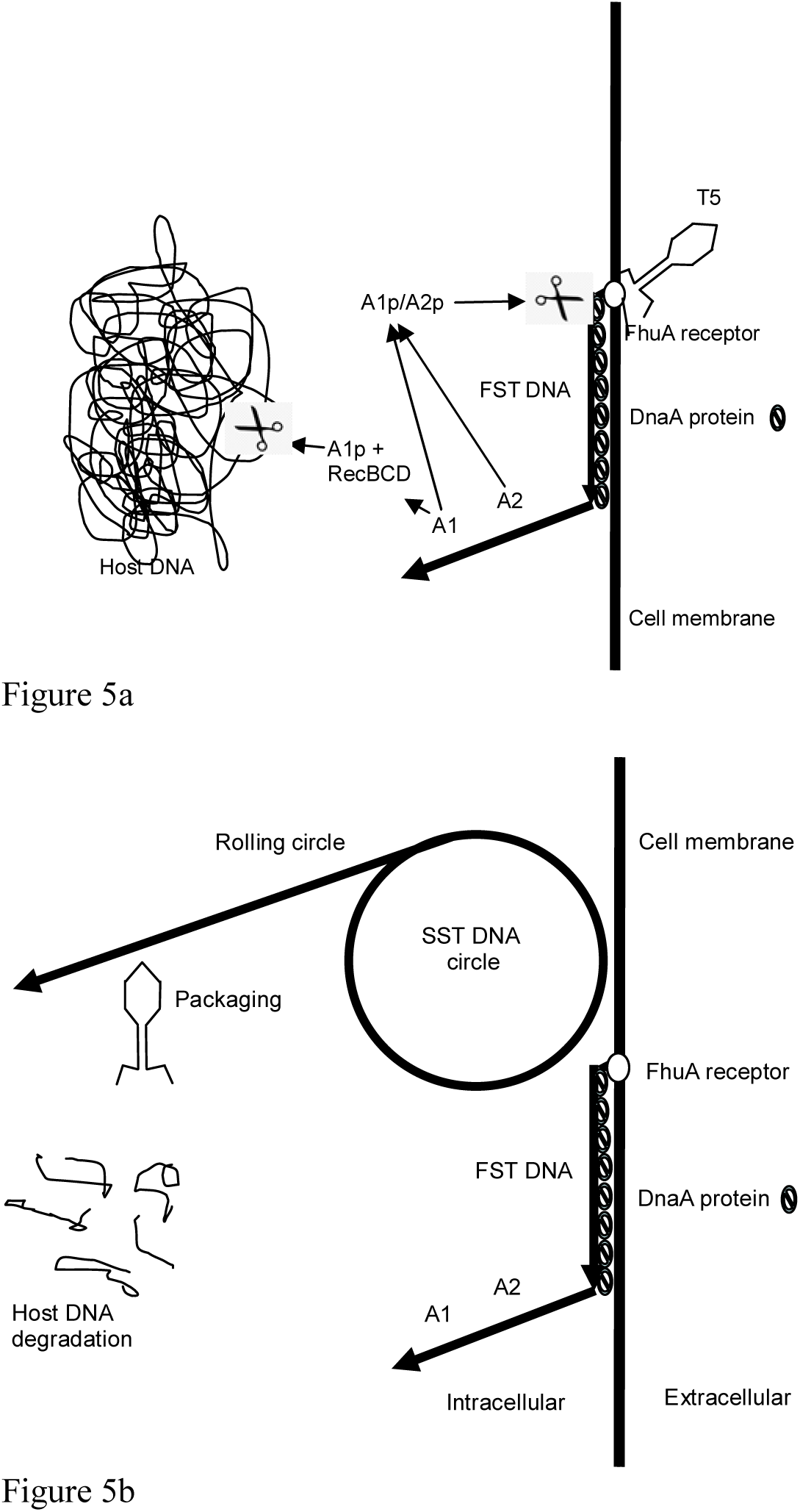
First and second step injection of the T5 genome. The vertical black line represents the membrane with the interior of the cell to the left and the outside to the right. Figure 5a: This figure is not drawn to scale. The T5 phage is bound to the FhuA receptor (open circle) and has injected the FST DNA (black arrow). The FST DNA is bound by its DnaA protein binding sites to the DnaA protein (barred circles) in the membrane. The A1 protein is a restriction endonuclease that cleaves the host DNA. Complete host DNA degradation is achieved by the host RecBCD nuclease. Figure 5b: This figure is nor drawn to scale and the rolling circle in is, in reality, about ten times the size of the FST DNA. The A1/A2 heterodimer endonuclease cleaves the FST DNA at a specific restriction site beyond the blockage caused by the DnaA protein binding sites and permits injection to continue. When the homologous restriction site on the RTR DNA arrives inside the cell, it is similarly cleaved. The two restriction sites join either by their cohesive ends or by site specific recombination catalyzed by the A1/A2 dimer. This results in circle formation that protects the phage DNA from degradation by RecBCD nuclease. This circle formation initiates DNA replication and eventually rolling circle formation which forms concatameric DNA. The concatameric DNA is then processed and packaged into phage particles by the terminase.

### 4.10 Predictions of the new restriction site model

Naturally, this new restriction site model of T5 two-step injection must be too simplistic since it assigns no role to the conserved multitude of repeats (inverted repeats and palindromes) in the *iss* region. It seems likely that these conserved structures are involved in the injection stop or re-start processes.

The restriction site model predicts the generation of a FST DNA fragment as a normal part of T5 infection (i.e. independent of shearing). Curiously, strong support for the restriction site model, comes from a 52 year old (1968) experiment by Lanni (2). In one control experiment she verified the DNA from cells infected with T5 but neither blended nor treated with cloramphenicol. As expected high molecular weight phage T5 DNA was present. However, a fragment corresponding in size to FST DNA Figure 6 was also clearly present. The authors did not attempt to assign a biological origin of this FST fragment, but instead suggested DNA breakage during extraction (47) leaving unclear why breakage of the large T5 DNA molecule would generate a fragment of FST size. The presence of this FST fragment demonstrates that the FST fragment does not only derive from blending (since no blending was used) but is a natural part of the infection process. This result is incompatible with the classical model whereby the T5 DNA molecule is injected intact. In contrast, the presence of this FST fragment is a prediction of the restriction enzyme model outlined above.

**Figure 6.**
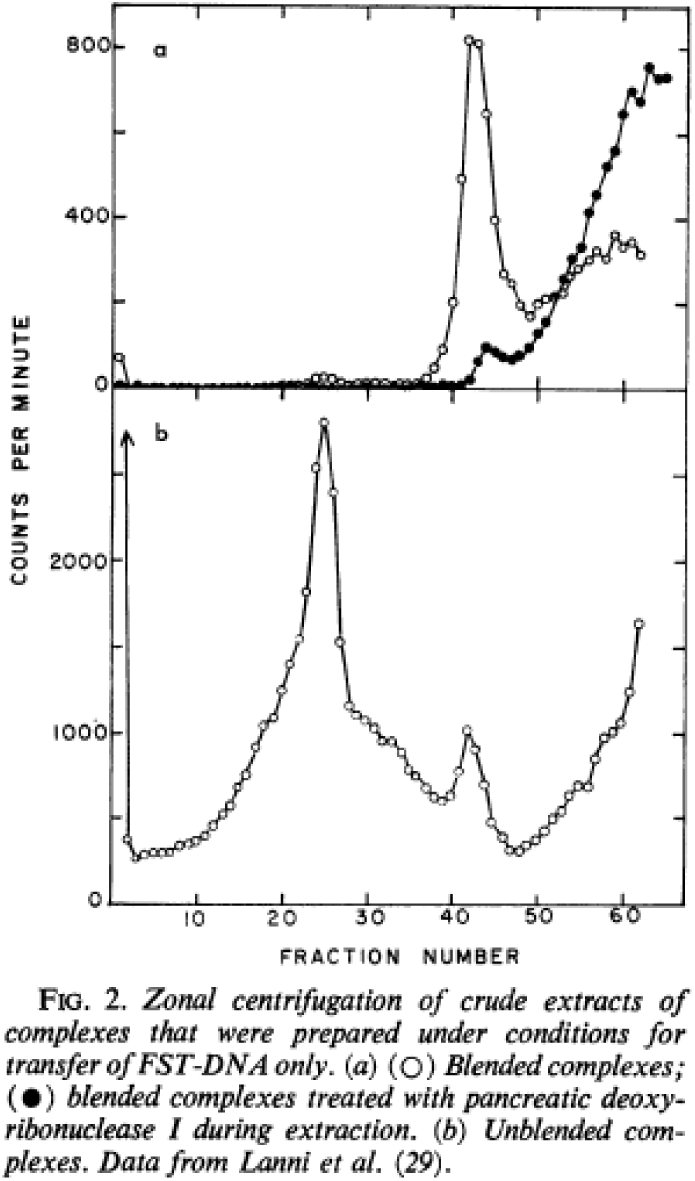
This figure is borrowed from the review by Lanni, 1968 (2). Figure la shows the classical demonstration of FST DNA, obtained by blending cloramphenicol treated T5-infected cells. Figure 2b shows the same infected cells but without either blending or cloramphenicol treatment. It should be noted that a band corresponding in size to the FST is present in untreated T5 infected cells.

The new model takes into account the nuclease properties of the A1 protein as well as the presence of precisely orientated DnaA boxes within the *iss* region. It provides a radically different viewpoint of two-step injection, and, unlike the Lanni model, it makes predictions that can be tested. Using this insight, it should be possible to test for restriction endonuclease activity in the early stages of T5 infection. The additional predictions that A2 modified A1 restriction endonuclease attacks restriction sites in each terminal repeat and that it may be involved in circularization, could also be tested experimentally.

It would be interesting to investigate whether T5 infection of a thermo-sensitive DnaA mutant, at the non-permissive temperature, would give a two-step injection or whether all of the DNA would enter the cell in one singe step; which would probably be lethal for both the phage and the host.

Finally, this report illustrates the difficulty of using genetics to investigate a cis-acting essential region of DNA where mutation would result in a lethal phenotype. A possible solution is to use a heterologous system. For example, it may be possible to clone the *iss* region into a small mobilizable plasmid vector (such as pJRD215 (48)). If so, perhaps the *iss* region would prevent conjugal transfer of the plasmid to another host and this may be a way to select mutants in the *iss* region. Such a possibility may open up the *iss* region to a new kind of genetic analysis.

## Notes

https://www.biorxiv.org/content/10.1101/866236v1

### Competing Interest Statement

The authors have declared no competing interest.

### Summary of Updates

Addition of new data and Supplemental files and Figure 5 have been added. References added

